# Heterogeneity of CD8αα intraepithelial lymphocytes is transcriptionally conserved between TCRαβ and TCRγδ cells

**DOI:** 10.1101/2025.05.20.655135

**Authors:** Kaito A. Hioki, Xueting Liang, Adam C. Lynch, Ravi Ranjan, Elena L. Pobezinskaya, Leonid A. Pobezinsky

## Abstract

Intestinal intraepithelial lymphocytes (IELs) are a versatile population of immune cells with both effector and regulatory roles in gut immunity. Although this functional diversity is thought to arise from distinct IEL subpopulations, the heterogeneity of TCRαβ^+^ and TCRγδ^+^ IELs have not been well-characterized. Using scRNAseq, we identified CD8αα^+^ T cell subsets with memory-like (*Tcf7*⁺) and effector-like (*Prdm1*⁺) profiles in both TCRαβ^+^ and TCRγδ^+^ IELs. Using CD160 and CD122 as markers of memory-like and effector-like cells, respectively, we found that while effector-like cells dominated the small intestine, memory-like IELs were more prevalent in the large intestine, suggesting a functional specialization of immune responses along the gut. Further transcriptional analysis revealed shared profiles between TCRαβ^+^ and TCRγδ^+^ small intestinal IEL subsets, suggesting conserved functional roles across these populations. Finally, our analysis indicated that TCRαβ^+^ memory-like IELs arise from *Tcf7⁺* double-negative (DN) precursors, and that effector-like IELs subsequently differentiate from the memory-like population. In contrast, TCRγδ^+^ IELs appear to originate from two distinct precursor populations, one expressing *Tcf7* and the other *Zeb2*, indicating the presence of parallel developmental pathways within this lineage. Overall, our findings reveal that both TCRαβ^+^ and TCRγδ^+^ cells contain memory-like and effector-like subsets, which may contribute to the functional heterogeneity of IELs.

## Introduction

The intestinal epithelium is populated by intraepithelial lymphocytes (IELs), a diverse group of T cells that play a critical role in maintaining gut health through several mechanisms. These include effector functions via the secretion of inflammatory cytokines or antimicrobial peptides, cytotoxic activity through perforin and granzymes, regulatory functions by tolerating food antigens or commensal microbes, and even promoting epithelial barrier repair after inflammation ^1–5^. It is hypothesized that various IEL subpopulations are responsible for these different roles. These subpopulations have been categorized as “induced IELs,” and “natural IELs”, corresponding to conventional tissue-resident T cells and unconventional T cells, respectively. Induced IELs develop from circulating conventional T cells and consist of the following subsets: TCRαβ^+^ CD4^+^, TCRαβ^+^ CD8αβ^+^, and TCRαβ^+^ CD4^+^ CD8αα^+^ (TCRαβ^+^ double-positive or TCRαβ^+^ DP). Natural IELs derive directly from thymic precursors and include TCRαβ^+^ CD4^-^ CD8αβ^-^ (TCRαβ^+^ double-negative or TCRαβ^+^ DN), TCRαβ^+^ CD4^-^ CD8αβ^-^ CD8αα^+^ (TCRαβ^+^ CD8αα^+^), TCRγδ^+^ CD4^-^ CD8α^-^ Cd8β^-^ (TCRγδ^+^ double-negative or TCRγδ^+^ DN), and TCRγδ^+^ CD4^-^ CD8αβ^-^ CD8αα^+^ (TCRγδ^+^ CD8αα^+^) cells ^2,4,6^.

Induced IELs are conventional TCRαβ^+^ cells that migrate to the intestine after encountering antigens and can perform effector functions resembling tissue resident memory cells. Induced TCRαβ^+^ CD8αβ^+^ cells account for 10-15% of all IELs ^7,8^. These cells are implicated in controlling viral infections and pathogenic bacteria by expressing high levels of effector markers, while producing limited amounts of inflammatory cytokines ^9–11^. Induced TCRαβ^+^ CD4^+^ IELs can exhibit both immune-suppressive and effector functions ^5,12,13^. Regulatory T cells (Tregs) and T-helper subsets produce IL-10 or TGF-βto suppress excessive inflammation in response to microbes ^3,14–16^. Furthermore, upon entering the intestine, some TCRαβ^+^ CD4^+^ IELs begin to express CD8αα in a microbiota-dependent manner and acquire cytotoxic features such as production of granzymes and IFN-β ^17–19^.

The function of natural IELs is incompletely understood, despite representing the majority of IELs. TCRαβ^+^ DN thymic emigrants begin expressing CD8αα upon entering the intestine, and acquire cytotoxic features such as granzymes, Fas-ligand, and NK cell receptors ^20,21^. However, these cells do not initiate proinflammatory responses against infections. Rather, they are believed to play a regulatory role in suppressing colitis and have a high activation threshold to self-antigens due to the absence of co-stimulation from CD8αβ ^22–24^. TCRγδ^+^ IELs display a more pronounced cytotoxic profile through their expression of proinflammatory cytokines and anti-microbial peptides. These cells have been shown to be critical in reducing tumor volume in colorectal cancer and lowering pathogenic microbial loads in colitis models ^3,25,26^. On the other hand, TCRγδ^+^ IELs also produce anti-inflammatory cytokines and profibrotic factors, which play important roles in intestinal epithelial repair and barrier maintenance ^1,27–30^. Given the polyfunctional nature of IELs, it is likely that higher resolution analyses could improve the characterization of IEL subsets.

Although various studies have implemented multicolor flow cytometry or high-throughput sequencing technologies to analyze IEL subpopulations, they have not sufficiently addressed the basis for the IEL functional heterogeneity. Here, we performed single-cell RNA sequencing (scRNAseq) on sorted TCRαβ^+^ and TCRγδ^+^ small intestinal IELs and identified CD8αα^+^ T cell subsets with distinct profiles. Both datasets revealed subsets with memory T cell-like characteristics expressing *Tcf7* (TCF1), and subsets with effector T cell-like characteristics expressing *Prdm1* (BLIMP1). Additionally, TCRγδ^+^ IELs contained a unique CD8αα^+^ cluster expressing *Zeb2*. Using surface markers CD160 and CD122 to distinguish memory-like and effector-like CD8αα^+^ IELs, respectively, we found that effector-like cells were more prevalent in the small intestine. This population of effector-like TCRαβ^+^ cells gradually decreased along the sections of the small intestine and was absent in the large intestine, where only a single population of CD122^low^ CD160^low^ cells was observed. In contrast, effector-like TCRγδ^+^ cells also decreased along the small intestine but were retained in the large intestine. These observations suggest that environmental factors may influence the distribution of memory-like and effector-like IELs, with similar effects observed for both TCRαβ^+^ and TCRγδ^+^ cells in the small intestine but varying impacts in the large intestine. Further analysis of our small intestinal scRNAseq dataset revealed significant transcriptional similarities between TCRαβ^+^ and TCRγδ^+^ cells. Finally, based on our data, we predicted the precursor-progeny relationships among different IEL subsets.

## Results

### TCRαβ^+^ and TCRγδ^+^ IELs consist of diverse groups of cells

While many studies have explored the heterogeneity of IELs using single-cell sequencing techniques, it has been a challenge to clearly distinguish between TCRαβ^+^ and TCRγδ^+^ cells at the transcriptional levels, in part due to the shared usage of Vα and V8 gene segments. To better characterize the populations of TCRαβ^+^ and TCRγδ^+^ IELs, we performed scRNAseq on FACS-sorted TCRαβ^+^ and TCRγδ^+^ IELs collected from the small intestine of healthy C57BL/6 mice. Uniform manifold approximation and projection (UMAP) analysis was performed separately for the TCRαβ^+^ and TCRγδ^+^ cells, revealing 16 and 17 clusters, respectively (**Fig. 1a,b**). Based on the expression level of *Cd4*, *Cd8a*, and *Cd8b1* coreceptor genes, we noticed that minor IEL populations defined in flow cytometry experiments (TCRαβ^+^ DN, TCRαβ^+^ DP, and TCRγδ^+^ DN; **Sup.** Fig. 1a) were not represented as distinct clusters (**Fig. 1c,d**), possibly due to the relatively small population sizes or stringent quality control filtering. Among the conventional TCRαβ^+^ cells, CD4^+^ cells comprised clusters 9, 14 and part of cluster 3, while CD8αβ^+^ cells consisted of clusters 2, 8, 12 and part of cluster 3. Surprisingly, our analysis of natural TCRαβ^+^ CD8αα^+^ cells revealed two distinct cluster groups: clusters 1, 4, 6, 15, and clusters 5, 7, 10 (**Fig. 1a,c**). Similarly, the TCRγδ^+^ CD8αα^+^ clusters formed diverging groups on the UMAP plot (**Fig. 1b,d**).

**Figure 1:**
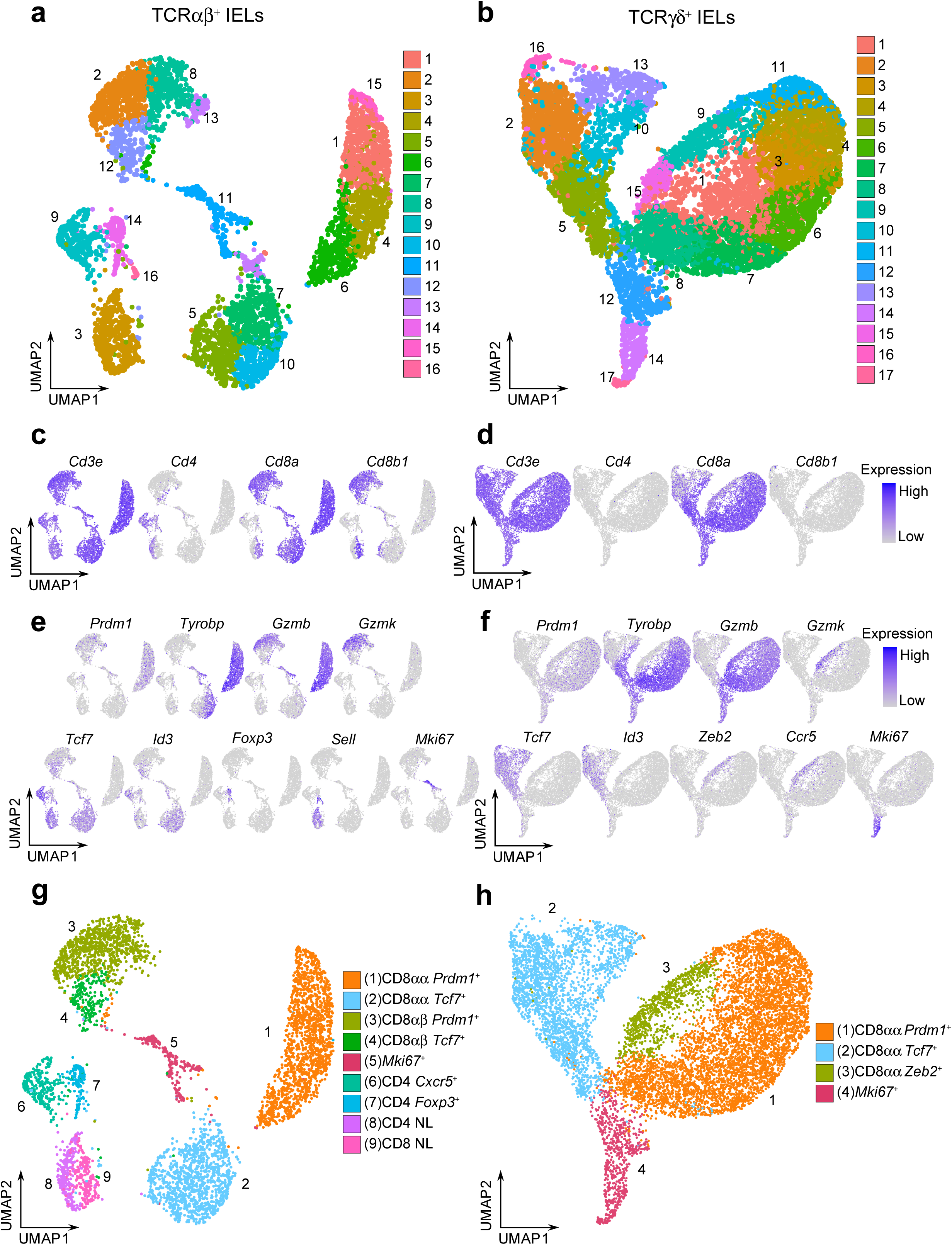
TCRαβ^+^ and TCRγδ^+^ IELs consist of diverse groups of cells. **a/b.** UMAP plots of sorted CD45^+^ TCRαβ^+^ TCRγδ^-^ IELs (a) or CD45^+^ TCRαβ^-^ TCRγδ^+^ IELs (b) after quality control filtering. **c/d.** Representative UMAP plots for the expression of *Cd3e* and coreceptor genes *Cd4*/*Cd8a*/*Cd8b1* for TCRαβ^+^ IELs (c) and TCRγδ^+^ IELs (d). **e/f.** Representative UMAP plots for the expression of marker genes descriptive of cluster group phenotypes for TCRαβ^+^ (e) and TCRγδ^+^ (f) datasets. **g/h.** Annotated UMAP plots of 4,845 TCRαβ^+^ IELs (g) and 10,418 TCRγδ^+^ IELs (h).

To define the phenotype of each cluster group, we summarized the expression level of marker genes (**Sup.** Fig. 1b,c). The conventional TCRαβ^+^ CD4^+^ clusters consist of T-follicular helper-like cells expressing *Cxcr5* and *Il21* (cluster 9), *Foxp3*^+^ Tregs (cluster 14), and naïve cells expressing *Sell*, *Ccr7*, and *Dapl1* (cluster 3) ^31–34^. The CD8αβ^+^ clusters 2 and 8 were enriched for effector-like T cell markers, including *Prdm1* (which encodes the transcription factor BLIMP1) and effector molecules *Gzmb* and *Gzmk* ^35^ (**Fig. 1e**). In contrast, cluster 12 was enriched for memory-like T cell markers, such as *Tcf7* (encoding the transcription factor TCF1) and *Id3*. CD8αβ^+^ cells were also detected in the naïve cell cluster (cluster 3). Interestingly, within the TCRαβ^+^ CD8αα^+^ clusters, we also observed two distinct groups: clusters 1, 4, 6 and 15 were enriched for effector-like markers, including *Prdm1*, *Tyrobp*, *Gzma* and *Gzmb* , whereas clusters 5, 7 and 10 were enriched for memory-like markers *Tcf7* and *Id3* ^35,36^ (**Fig. 1e**). A similar separation between effector-like and memory-like profiles was found among the TCRγδ^+^ cells (clusters 1, 3, 4, 6, 7, 8, 11 and clusters 2, 5, 10, 13, respectively; **Fig. 1f**).

Collectively, we annotated cluster groups according to the expression of enriched transcription factors: the effector-like cluster groups were labeled as “*Prdm1*^+^” clusters, and the memory-like cluster groups were labeled as “*Tcf7*^+^” clusters (**Fig. 1g, h**). Additionally, a unique effector-like population within the TCRγδ^+^ IELs characterized by *Zeb2*, *Gzmk*, and *Ccr5*, was labeled as “*Zeb2^+^*” cells (former clusters 9 and 15). Finally, cluster groups enriched in proliferation and cell cycle gene markers were annotated as “*Mki67*^+^” clusters (former TCRαβ^+^ cluster 11 and TCRγδ^+^ clusters 12, 14, 17). Clusters with poorly defined identities or nonspecific cluster localization (former TCRαβ^+^ clusters 13 and 16, and TCRγδ^+^ cluster 16) were excluded from further analysis. In our final annotation, we retained 4,845 TCRαβ^+^ IELs and 10,418 TCRγδ^+^ IELs.

CD8αα^+^ IEL subsets have distinct transcriptional features and preference in colonization pattern of the intestine TCRαβ^+^ and TCRγδ^+^ CD8αα^+^ IELs are considered “non-conventional” T cells, due to their unique developmental pathways, TCR-independent activity, and other innate-like features. These cells play versatile roles in gut immunity, including the secretion of antimicrobial peptides during infections, exhibiting cytotoxic potential, and supporting repair after inflammation, while also displaying higher activation thresholds for self-reactivity ^1,3,37,38^. Our observation of the distinct CD8αα^+^ subsets may provide insight into their functional diversity.

To better understand the differences between the CD8αα^+^ sub-clusters, we filtered out cells which expressed *Cd4* or *Cd8b1* coreceptor genes and re-clustered the populations (**Sup.** Fig. 2a, b). When confirming the expression of *Cd8a*, we noticed the re-clustered cells included those with no *Cd8a* expression in each of the annotated populations (**Sup.** Fig. 2c, d), The coreceptor negative cells contributed to 4.7% and 11.9% of the re-clustered TCRαβ^+^ and TCRγδ^+^ cells, respectively, and were localized to the edges of the UMAP plots (**Sup.** Fig. 2e, f). We speculated that these cells represent the population of DN IELs identified by flow cytometry experiments (**Sup.** Fig. 1a), which are known as precursors to CD8αα^+^ IELs ^13^. Accordingly, we performed the subsequent analyses separately for the *Cd8a*^+^ (CD8αα^+^) IELs and coreceptor negative DN IELs, with the cluster labels identified in **Figure 1**.

We first compared the transcriptional profiles of the CD8αα^+^ sub-clusters (**Sup. File**). In both the TCRαβ^+^ and TCRγδ^+^ datasets, *Prdm1*^+^ clusters were significantly enriched for effector T cell-like transcription factors including *Prdm1*, *Runx1*, *Runx2*, *Plek* and *Irf8*. In contrast, *Tcf7*^+^ clusters showed marked enrichment for memory-like transcription factors such as *Tcf7*, *Id3*, *Aff3*, *Bach2*, *Nfkb1* and *Satb1* (**Fig. 2a-d**). Additionally, several transcription factors displayed lineage-specific expression pattern. For example, *Foxo3*, *Jazf1*, *Zbtb16* and *Lef1* were differentially expressed among clusters within the TCRαβ^+^ lineage, whereas *Zmat4* and *Zfnx3* and *Elk4* were specific for TCRγδ^+^ cells. Interestingly, within the TCRγδ^+^ dataset, the *Zeb2*^+^ cluster exhibited a transcriptional profile intermediate between *Prdm1*^+^ and *Tcf7*^+^ clusters, while also expressing unique TFs such as *Zeb2*, *Bcl11b* and *Rora* (**Fig. 2d**). The CD8αα^+^ *Prdm1*^+^ and *Tcf7*^+^ clusters differentially expressed genes encoding surface proteins, with *Il2rb* (CD122) being enriched in the *Prdm1*^+^ clusters, and *Cd160* (CD160) predominantly expressed in *Tcf7*^+^ clusters (**Fig. 2e, f**). Both CD122 and CD160 have been previously studied for their roles in IEL maintenance, cytotoxic activity, or protective ability, but their heterogeneous expression within total IEL populations has not been fully clarified ^37,39–42^. To validate whether CD160 and CD122 can be used to distinguish memory-like and effector-like subsets of CD8αα^+^ IELs, we performed flow cytometry on small intestinal IELs stained for these surface markers. The analysis revealed two distinct populations, with approximately 20% of CD8αα^+^ IELs being CD122^int^CD160^+^ and 75% being CD122^hi^CD160^-^ (**Fig. 2g**). These results suggest that the effector-like and memory-like phenotypes are distinctly distributed among total IELs, with surface markers CD160 and CD122 providing a means to distinguish between these functional subsets.

**Figure 2:**
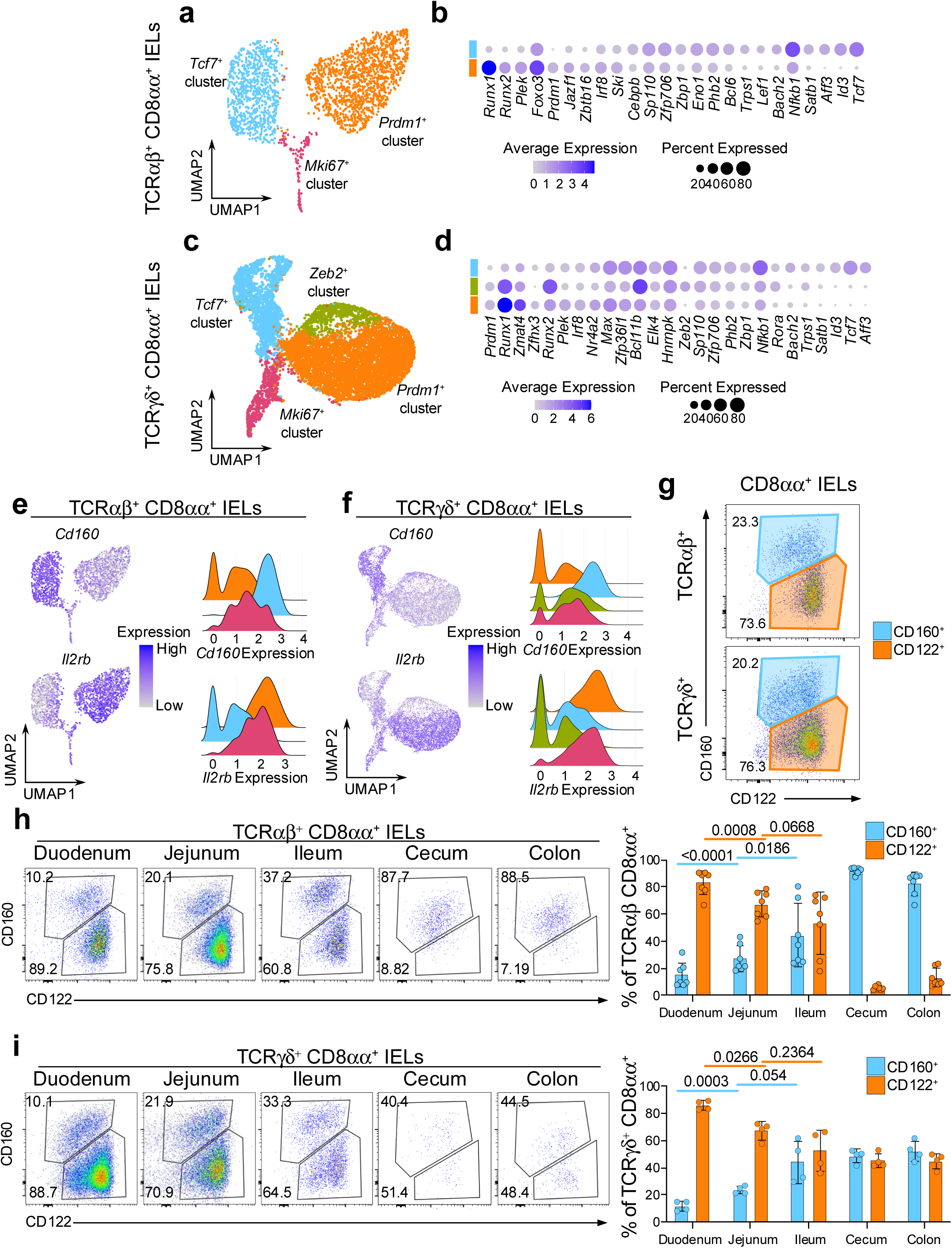
CD8αα^+^ IEL subsets have distinct transcriptional features and colonization pattern of the intestine. **a.** UMAP plot of re-clustered TCRαβ^+^ CD8αα^+^ cells. **b.** Differentially expressed transcription factors between the (non-*Mki67*^+^) CD8αα^+^ cluster groups in TCRαβ^+^ CD8αα^+^ cells. **c.** UMAP plot of re-clustered TCRγδ^+^ CD8αα^+^ cells. **d.** Differentially expressed transcription factors between the (non-*Mki67*^+^) CD8αα^+^ cluster groups in TCRγδ^+^ CD8αα^+^ cells. **e/f.** Expression of differentially expressed surface protein-coding genes represented as UMAP plots and ridge plots for TCRαβ^+^ CD8αα^+^ cells (e) and TCRγδ^+^ CD8αα^+^ cells (f). **g.** Representative flow cytometry plots of CD122 and CD160 expression among TCRαβ^+^ CD8αα^+^ cells and TCRγδ^+^ CD8αα^+^ cells from the small intestine. **h/i.** Representative flow cytometry plots and quantification of CD122 and CD160 expression along five sections of the intestine for TCRαβ^+^ CD8αα^+^ cells (h) and TCRγδ^+^ CD8αα^+^ cells (i). Data were analyzed by Paired T-test (h,i).

The maintenance and activity of IELs can be modulated by both diet and gut microbiota, which vary along the intestinal tract ^17,18,43–48^. These variations may shape the distribution of IEL subsets to ensure appropriate immune responses in different regions of the intestine. Therefore, we next examined the composition of CD8αα^+^ IELs along five different sections of the intestine. In both TCRαβ^+^ and TCRγδ^+^ CD8αα^+^ IELs, the proportion of CD160^+^ cells gradually increased from 10% to 30% along the small intestine (duodenum, jejunum, ileum), while the proportion of CD122^hi^ cells decreased reciprocally (**Fig. 2h, i**). However, the distribution of these populations shifted drastically in the large intestine. Among TCRαβ^+^ IELs, CD122^+^ cells were nearly absent in cecum and colon, leaving one major population of CD122^int^CD160^int^ cells (**Fig. 2h**). In contrast, despite collecting very few of TCRγδ^+^ IELs in the large intestine, an equal proportion of CD122^hi^CD160^-^ and CD122^int^CD160^int^ populations were observed (**Fig. 2i**).

To confirm whether the CD160^+^ population in all five sections of the intestine correspond to the *Tcf7*^+^ cluster IELs, we used *Tcf7*-GFP mice, where *Tcf7*-expressing cells are concurrently labeled by GFP ^49^. Indeed, the CD8αα^+^ CD160^+^ IELs expressed higher levels of GFP compared to CD8αα^+^ CD122^+^ cells in most tissue sections, for both TCRαβ^+^ and TCRγδ^+^ subsets (**Sup.** Fig. 3a). These observations suggest that some yet unknown factors influence the abundance of CD8αα^+^ IELs along the intestine, with similar prevalence of effector-like IELs in TCRαβ^+^ and TCRγδ^+^ cells in the small intestine but favoring the colonization of memory-like TCRαβ^+^ CD8αα^+^ IELs in the large intestine.

### DN IEL subsets have distinct transcriptional features and no preference in colonization along the intestine

Using our dataset of coreceptor-negative DN IELs, we next examined whether the transcriptional differences between the *Prdm1*^+^ and *Tcf7*^+^ clusters could also be observed at this stage. Gene expression analysis across both TCRαβ^+^ and TCRγδ^+^ lineages revealed that DN subsets share many transcription factors with their corresponding CD8αα^+^ counterparts. Specifically, both *Tcf7*^+^ DN and CD8αα^+^ IELs expressed *Tcf7* and *Zfp706,* while *Prdm1^+^* subsets shared expression of *Prdm1* and *Runx1* (**Fig. 2a-d and 3a-d**). Furthermore, we found a strong overlap in transcription factor expression between DN and CD8αα^+^ subsets within the TCRγδ^+^ lineage (**Fig. 2a-d and 3a-d**). For example, *Zmat4* was highly enriched in the *Prdm1*^+^ cluster; *Zeb2*, *Runx2*, *Stat1*, and *Bcl11b* were enriched in the *Zeb2^+^* cluster; and *Tcf7*, *Id3*, *Aff3*, *Satb1*, *Lef1*, and *Nfkb1* were enriched in the *Tcf7*^+^ cluster (**Fig. 2d and 3d**). In addition, we identified several genes, which were specific to *Tcf7^+^* DN subsets, including TFs associated with the cell cycle such as *Hnrnpk*, and *Ybx1* ^50–52^ (**Fig. 3a-d**), along with increased expression of ribosomal genes, a pattern not observed among the CD8αα^+^ subclusters (**Sup. file**). These findings suggest that DN cells represent distinct populations that nonetheless relate to CD8αα^+^ cells, supporting the possibility of developmental continuum between these subsets.

**Figure 3:**
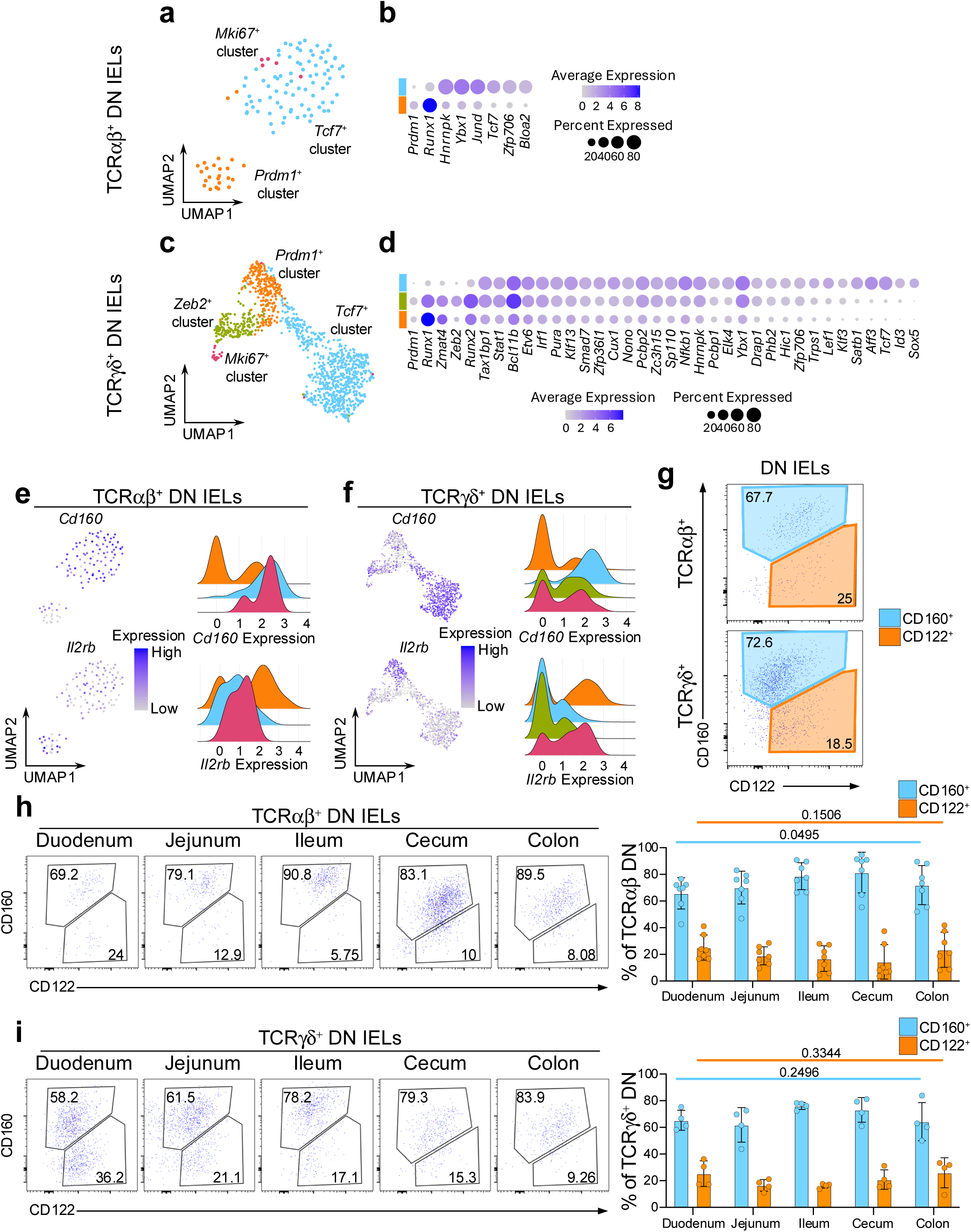
DN IEL subsets have distinct transcriptional features and static colonization pattern of the intestine. **a.** UMAP plot of re-clustered TCRαβ^+^ DN cells. **b.** Differentially expressed transcription factors between the (non-*Mki67*^+^) DN cluster groups in TCRαβ^+^ DN cells. **c.** UMAP plot of re-clustered TCRγδ^+^ DN cells. **d.** Differentially expressed transcription factors between the (non-*Mki67*^+^) DN cluster groups in TCRγδ^+^ DN cells. **e/f.** Expression of differentially expressed surface protein-coding genes represented as UMAP plots and ridge plots for TCRαβ^+^ DN cells (e) and TCRγδ^+^ DN cells (f). **g.** Representative flow cytometry plots of CD122 and CD160 expression among TCRαβ^+^ DN cells and TCRγδ^+^ DN cells from the small intestine. **h/i.** Representative flow cytometry plots and quantification of CD122 and CD160 expression along five sections of the intestine for TCRαβ^+^ DN cells (h) and TCRγδ^+^ DN cells (i). Data were analyzed by one-way ANOVA (h,i).

We also examined the expression level of surface protein coding genes which were differentially expressed between CD8αα^+^ subclusters. Similarly to the CD8αα^+^ IELs, the *Tcf7*^+^ DN clusters had greater mRNA expression of *Cd160*, while the *Prdm1*^+^ DN clusters showed higher expression of *Il2rb* (**Fig. 3e, f**). These markers also distinguished DN IELs at the protein level by surface expression. However, in contrast to the CD8αα^+^ IELs, which contained a greater proportion of CD122^hi^ cells (**Fig. 2g**), approximately 70% of DN IELs were CD160^+^ and 20% were CD122^hi^ (**Fig. 3g**). Based on *Tcf7*-GFP reporter expression, we observed a clear trend (although not statistically significant) indicating that CD160+ DN cells expressed higher levels of *Tcf7* (**Sup.** Fig. 3b). These findings align with the predicted distribution from the sequencing data (**Sup.** Fig. 2e, f). Of note, unlike natural IELs, induced IELs were more uniform and did not clearly segregate into memory-like and effector-like phenotype based on expression of CD122 and CD160 (**Sup.** Fig. 3c). An exception was observed for TCRαβ^+^ CD8αα^+^ CD4^+^ (DP) cells, where the majority of cells displayed an effector-like CD160^-^CD122^int^ phenotype.

Interestingly, unlike CD8αα^+^ IELs, the proportion of CD122^+^ and CD160^+^ cells along the intestine remained stable among both TCRαβ^+^ and TCRγδ^+^ DN IELs (**Fig. 3h, i**). This suggests that CD8αα^+^ IEL subsets are uniquely sensitive to yet unidentified microenvironmental cues that may drive their “gradient-like” distribution along the small intestine. Notably, this unique sensitivity appears to be specific to the small intestine, as both the CD8αα^+^ and DN IELs in the large intestine (cecum and colon) are predominantly comprised of CD122^low^CD160^int^ cells (**Fig. 2h, i, and Fig. 3h, i**).

### The transcriptional profiles of TCRαβ^+^ cells are similar to TCRγδ^+^ cells

The characteristics of CD8αα^+^ IELs have been suggested to vary between TCRαβ^+^ and TCRγδ^+^ cells, based on functional studies ^53^. However, the clustering profiles of the small intestinal CD8αα^+^ IELs appeared strikingly similar between the TCRαβ^+^ and TCRγδ^+^ datasets (**Fig. 2, 3**), suggesting the presence of subsets with shared phenotypes across both cell lineages. To further investigate this, we examined the similarity of IEL subsets between the two datasets.

First, we identified the gene signatures representing the CD8αα^+^ *Prdm1*^+^ (effector-like) and CD8αα^+^ *Tcf7*^+^ (memory-like) clusters in the total TCRαβ^+^ dataset. We calculated the differentially expressed genes comparing each subcluster to the rest of the TCRαβ^+^ cells and sorted the genes by their statistical significance (**Fig. 4a**). The top 10 representative genes of the TCRαβ^+^ CD8αα^+^ *Prdm1*^+^ cluster included genes associated with CD8 T cell activation such as *Fcer1g*, *Cd244a* (encoding 2B4), and *Clnk* (a SLP-76 member), or genes associated with cell-cell contacts such as *Fgl2*, *Frmd5*, and *Osbpl3* ^54–59^. The top 10 genes representing the TCRαβ^+^ CD8αα^+^ *Tcf7*^+^ cluster included genes associated with memory or stem-like T cells such as *Id3*, *Batf3*, *Kit*, and *AW112010* ^33^, and genes associated with intestinal resident T cells such as *Cd160* and *Xcl1*. Next, we projected the signature of each TCRαβ^+^ CD8αα^+^ subcluster onto the TCRγδ^+^ dataset to evaluate whether similarly annotated clusters showed similar enrichment patterns. Indeed, the TCRαβ^+^ CD8αα^+^ *Prdm1*^+^ signature was significantly enriched in the TCRγδ^+^ CD8αα^+^ *Prdm1*^+^ clusters (**Fig. 4b**), indicating strong similarities between effector-like clusters of the TCRαβ^+^ and TCRγδ^+^ datasets. In parallel, the TCRαβ^+^ CD8αα^+^ *Tcf7*^+^ signature was highly enriched in the TCRγδ^+^ CD8αα^+^ *Tcf7*^+^ clusters, highlighting a similar correspondence between memory-like populations. Together, these findings underscore striking parallels between IEL subsets across the TCRαβ^+^ and TCRγδ^+^ lineages.

**Figure 4:**
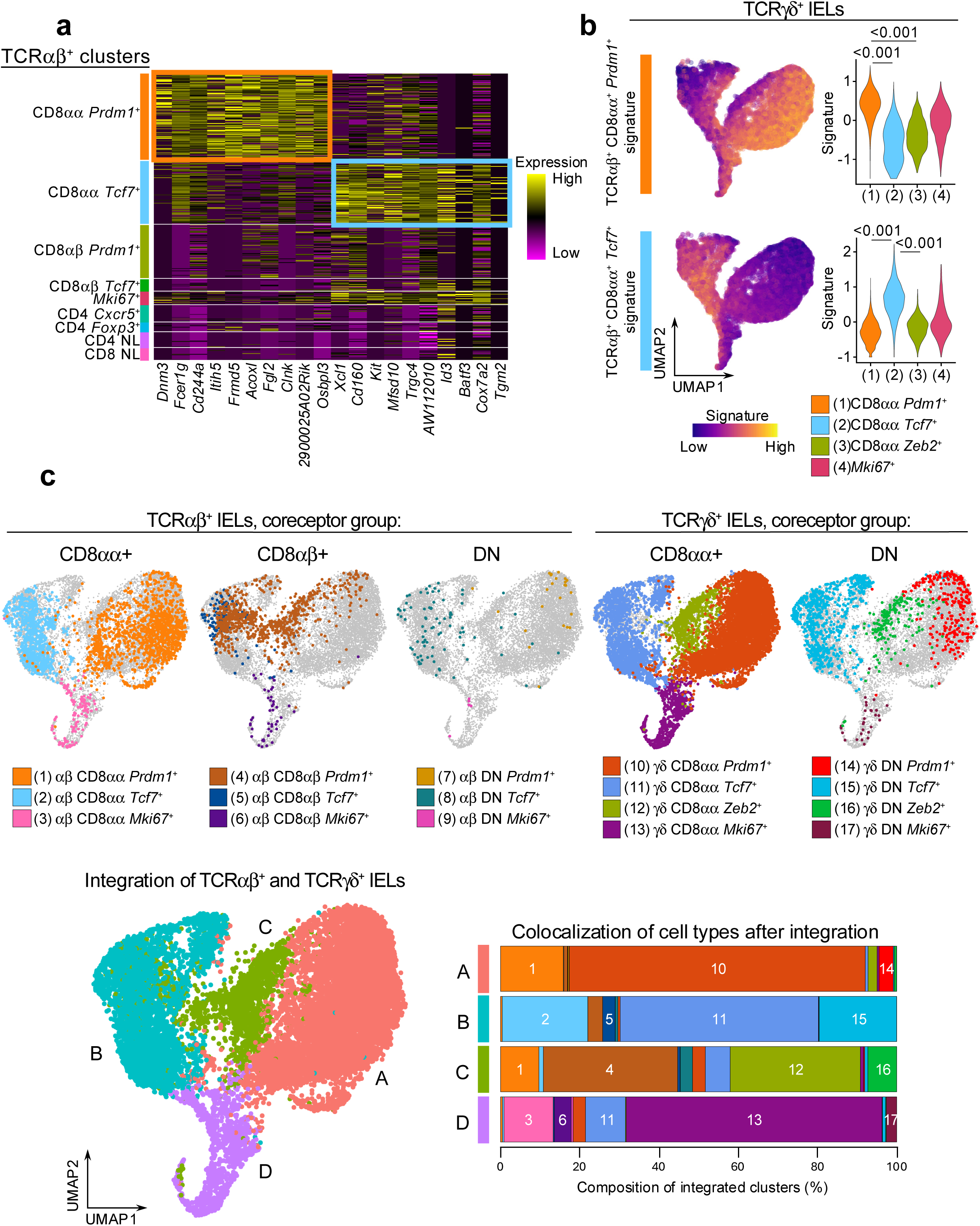
The transcriptional profiles of TCRαβ^+^ cells are similar to TCRγδ^+^ cells. **a.** Heatmap of top 10 marker genes for the annotated TCRαβ^+^ IEL subclusters sorted by statistical significance. **b.** Expression of CD8αα^+^ cluster marker genes in (a) among total TCRγδ^+^ cells. Signature scores were compared by Mann-Whitney U test. **c.** UMAP plot generated after integration of TCRαβ^+^ and TCRγδ^+^ datasets with original cluster identities overlaid (top). Integrated clusters (bottom left) and quantification of cluster composition by individual cluster groups represented as a bar plot (bottom right).

Although these findings validate the overlap between corresponding clusters, it is possible that additional parallels exist between the two datasets that were not detected using a signature projection approach. For example, neither of the signatures showed enrichment in the TCRγδ^+^ CD8αα^+^ *Zeb2^+^* cluster. To further evaluate the similarities between TCRαβ^+^ and TCRγδ^+^ IEL clusters, we applied a second approach, where we integrated populations from the two datasets and re-clustered them with low resolution (**Fig. 4c**). In addition to the CD8αα^+^ and DN IELs analyzed in **Figure 2 and 3**, we also included the TCRαβ^+^ CD8αβ^+^ clusters as they represent a major portion of TCRαβ^+^ IELs (**Sup.** Fig. 1a). As expected, the integrated UMAP analysis showed that the “*Tcf7*^+^”, “*Prdm1*^+^” and “*Mki67*^+^” clusters from TCRαβ^+^ and TCRγδ^+^ cells grouped together. Similar to the previous observation (**Fig. 2d**), the TCRγδ^+^CD8αα^+^ *Zeb2^+^* cluster grouped together with a portion of the TCRαβ^+^ CD8αβ^+^ *Prdm1*^+^ cells, despite these clusters representing separate lineages of IELs. Altogether, our results demonstrate that many of the TCRαβ^+^ and TCRγδ^+^ cells have similar transcriptional profiles, suggesting a potential convergence of their functional roles in intestinal epithelial immunity.

### Prediction of precursor-progeny relationship between memory-like and effector-like IELs

Next, we investigated precursor-progeny relationships among natural IEL subsets across TCRαβ^+^ and TCRγδ^+^ lineages. Given that conventional memory T cells can differentiate into effector T cells, we asked whether a similar relationship exists between memory-like IEL clusters and effector-like IEL clusters, and whether such transitions could be inferred from our scRNAseq datasets. Since thymic IEL-precursors are phenotypically DN cells, we also explored the possibility of a developmental trajectory from DN cells to CD8αα^+^ IELs. Upon re-clustering of TCRαβ^+^ CD8αα^+^ and DN cells, we observed that only the *Tcf7*^+^ cluster contained a substantial proportion of DN cells (**Sup.** Fig. 2c, e), suggesting a single precursor population for TCRαβ^+^CD8αα^+^ IELs. In contrast, the TCRγδ^+^ dataset contained two clusters, *Tcf7*^+^ and *Zeb2*^+^, that were both enriched for DN cells (**Sup.** Fig. 2d, f), indicating the potential existence of two distinct precursor populations for TCRγδ^+^CD8αα^+^ IELs. To test whether these hypotheses are supported by computational predictions, we performed RNA velocity analysis on combined datasets of TCRαβ^+^ CD8αα^+^ and DN cells, as well as TCRγδ^+^ CD8αα^+^ and DN cells (**Fig. 5a, b**).

**Figure 5:**
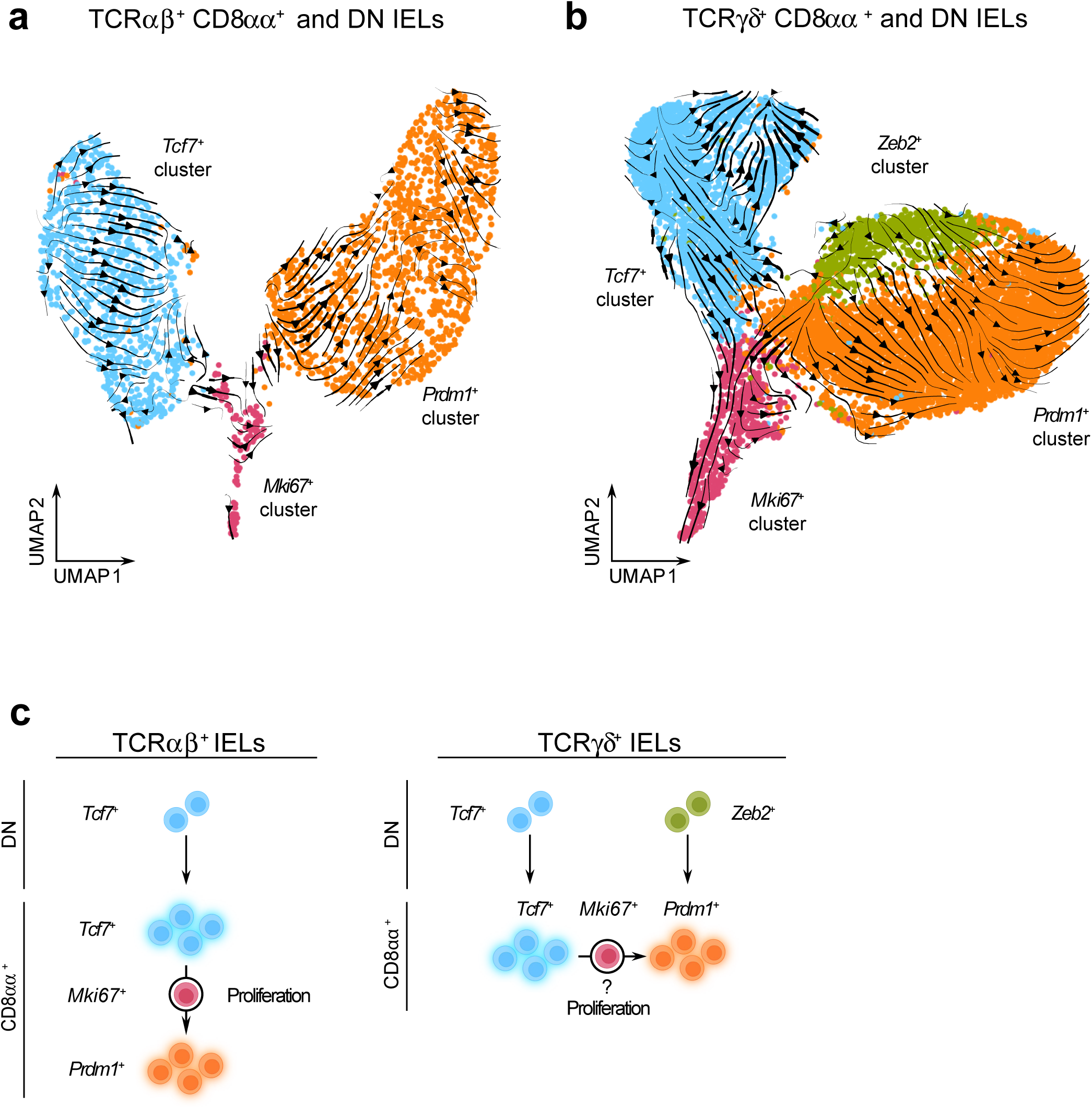
Memory-like and Effector-like CD8αα^+^ IELs may represent separate IEL lineages. **a/b.** UMAP plots with the overlay of RNA velocity trajectory predictions among re-clustered TCRαβ^+^ CD8αα^+^ and DN cells (a) and re-clustered TCRγδ^+^ CD8αα^+^ and DN cells (b). **c.** Trajectory models between IEL clusters predicted by RNA velocity analysis.

The TCRαβ^+^ dataset contained distinct trajectories within both the *Tcf7*^+^ and *Prdm1*^+^ clusters. Notably, *Tcf7*^+^ DN cells were positioned at the beginning of the trajectory, clustering together at the earliest point, and therefore appeared as sole precursors to more differentiated *Tcf7*^+^ CD8αα^+^ cells (**Sup.** Fig. 2e, **and Fig. 5a**). Along this trajectory, cells within the *Tcf7*^+^ cluster progressively downregulated the expression of transcription factor *Lef1* and the activating receptors *Klrk1* and *Klrc2,* while upregulating the memory- associated transcription factor *Batf3* ^60,61^. In contrast, the trajectory within the *Prdm1*^+^ cluster started near the *Ki67*^+^ population and was characterized by increased expression of *Zbtb16*, which encodes the innate-like T cell transcription factor PLZF ^62,63^ (**Sup.** Fig. 4a, b, c). These findings indicate the presence of phenotypic gradients within TCRαβ^+^ IEL compartment. Moreover, the directionality of the RNA velocity vectors suggested a developmental transition from the *Tcf7*^+^ cluster to *Prdm1*^+^ cluster, with the *Mki67*^+^ population acting as an intermediate proliferating stage. Thus, our results support the notion that DN *Tcf7^+^*cells function as precursors to natural TCRαβ^+^ CD8αα^+^ IELs (**Fig. 5c**).

The TCRγδ^+^ dataset also revealed lineage trajectories originating from DN cells (**Sup.** Fig. 2f, **and Fig. 5b**). Within the *Tcf7*^+^ cluster, two potential trajectory directions were observed. The first followed a linear progression from DN cells to CD8αα^+^ cells, marked by upregulation of memory-associated genes *Xcl1* and *Batf3,* and downregulation of genes involved in early T cell development, including *Rgs10* and *Eya2* ^64,65^ (**Sup.** Fig. 4d, e, f). The second trajectory, directed toward cluster 9 (**Sup.** Fig. 4d) and included contributions from both the DN and CD8αα^+^ cells, but lacked clearly distinguishable features. In contrast, the trajectory within the effector-like clusters appeared to originate from the DN cells in the *Zeb2^+^* cluster and progress toward CD8αα^+^ cells in the *Prdm1*^+^ cluster, and was characterized by high expression of effector-associated genes such as *Tyrobp*, *Fcgr3*, and *Ccrl2* (**Sup.** Fig. 4d, e, f). We also cannot rule out a possibility that similar to what was observed in the TCRαβ^+^ dataset, a transition from the *Tcf7*^+^ cluster to the *Prdm1*^+^ cluster may occur in TCRγδ^+^ lineage as well, via an intermediate *Mki67*^+^ proliferative population. Collectively, our data support the hypothesis that TCRγδ^+^ IELs arise from two independent precursor subsets (**Fig. 5c**).

We also considered an alternative hypothesis that the memory-like and effector-like IELs arise from separate precursors with their fates predetermined prior to maturation in the intestine. In mice, TCRαβ^+^ CD8αα^+^ IELs are known to develop from two thymic precursor populations that seed the intestine at different developmental stages and express oligoclonal TCR variable chains ^21,66–68^. To test whether memory-like or effector-like IELs exhibit distinct TCR Vα-chain usage, we measured the expression of Vα2 and Vα3.2 on the surface of TCRαβ^+^ CD8αα^+^ IELs from five sections of the intestine. However, we did not observe consistent enrichment of either Vα-chain among memory-like or effector-like subsets (**Sup.** Fig. 5), arguing against the idea that these populations originated from separate precursors. Altogether, these findings support a model in which memory-like and effector-like CD8αα^+^ IELs represent cells at different stages of shared differentiation pathways, enabling a spectrum of immune responses across the intestinal mucosa.

## Discussion

The current understanding for the heterogeneity of IELs has been suggested by a wide array of functional phenotypes attributed to surface proteins, transcription factor activity, or even developmental programs. In order to better define IEL subpopulations, single-cell sequencing experiments have been performed on total T cells; however, the distinction between TCRαβ^+^ and TCRγδ^+^ cells is difficult to make based on mRNA levels of TCR genes ^20,69,70^. To overcome this challenge, we sorted TCRαβ^+^ and TCRγδ^+^ small intestinal IELs and performed single-cell RNAseq. Clustering analyses for TCRαβ^+^ cells and TCRγδ^+^ cells revealed distinct populations of natural IELs based on marker genes. One population resembled memory-T cells, as indicated by the expression of *Tcf7* and *Id3*, and the other resembled effector-T cells, characterized by the expression of *Gzma*, *Gzmb*, *Tyrobp*, and *Prdm1*. The distinction between these memory-like and effector-like cells was present in both the coreceptor negative DN IELs and the CD8αα^+^ IELs. Our findings complement previous studies, which note similar memory-like and effector-like clusters in non-TCR-sorted IEL datasets ^26,35,36,71,72^. Although we have not examined the functional capabilities of effector-like and memory-like CD8αα^+^ IELs, others have highlighted the roles for these subsets. For example, effector-like TCRγδ^+^ cells in the colon demonstrated anti-tumor activity which was suppressed by *Tcf7* expression ^73^. Additionally, the cytotoxic potential of effector-like IELs has been shown in DSS-induced colitis, where tissue damage was reduced in *Ikzf3* KO mice with decreased amounts of effector-like IELs ^26^. Further studies will be necessary to elucidate the role of memory-like subsets, including their potential stem-like properties and anti-inflammatory functions in intestinal tissue-specific models. Interestingly, the similarly annotated clusters in our TCRαβ^+^ and TCRγδ^+^ datasets exhibited comparable transcriptional profiles. This suggests an overlap of cell phenotypes or functional contributions to intestinal immunity between TCRαβ^+^ and TCRγδ^+^ cells, despite the conventional view that IELs from these two lineages exhibit separate anti-inflammatory, regenerative, or cytotoxic roles ^1,3,5^. Our data suggests the functional heterogeneity of IELs can be further defined across the subsets of TCRαβ^+^ and TCRγδ^+^ cells.

Within our TCRγδ^+^ dataset, we also identified a novel cluster of effector-like CD8αα^+^ cells with high expression of *Zeb2, Ccr5* and *Gzmk*. *Zeb2* is linked to the terminal differentiation of effector CD8 T cells in infection models, whereas *Ccr5* is associated with infection induced migration to the gut mucosa ^74–76^. Furthermore, the transcriptional profile of this cluster overlapped with that of the effector-like TCRαβ^+^ CD8αβ^+^ cluster, suggesting that it may have been overlooked in previous studies that did not separate the TCRαβ^+^ and TCRγδ^+^ cells. This TCRγδ^+^ *Zeb2*^+^ subset may represent a transitional population of cells with a differentiation program resembling that of induced TCRαβ^+^ IELs. Further work is necessary to better define the phenotype and function of this IEL population.

We observed that the proportions of CD160^+^ and CD122^+^ cells among the CD8αα^+^ population are comparable between TCRαβ^+^ and TCRγδ^+^ IELs, suggesting that the distribution of these cell types in intestinal tissue may be influenced by environmental factors, such as dietary food and microbial antigens. While many studies have highlighted the importance of these factors in the development of major IEL subsets, few have inspected their impact on the more specific subsets of CD8αα^+^ cells ^35,73^. Yakou et al. observed a partial influence of microbiota, where germ free mice had a reduced number of TCF1^+^ memory-like cells in the colon but had no reduction of TCF1^+^ cells in the small intestine ^73^. Wang et al. demonstrated a partial impact of altered diet, where mice fed a high-fat high-sucrose “Western” diet reduced the amount of effector-like CD8αα^+^ IELs but increased the abundance of memory-like CD8αα^+^ IELs in the small intestine, with no difference in cell viability between the two populations ^35^.

We also identified transcriptionally distinct DN IEL populations. In both TCRαβ^+^ and TCRγδ^+^ lineages, the majority of DN cells belonged to the *Tcf7^+^* cluster. Additionally, the *Zeb2*^+^ cluster within TCRγδ^+^ IELs contained a unique subset of DN cells. Unlike CD8αα^+^ IELs, where the ratio between *Tcf7^+^* and *Prdm1*^+^ populations changes along the intestinal tract, the dominance of *Tcf7^+^*DN IELs remained consistent throughout the gut. This suggests that DN IELs are less responsive to local microenvironmental cues compared to their CD8αα^+^ counterparts. Additional studies exploring the relationship between the proportions of IEL subsets, microbial communities, and diet will be needed to clarify the heterogenous distribution of these cells across different parts of the intestine, and to determine the factors that render the CD8αα^+^ IELs sensitive to their dynamic tissue environments.

Through our analysis of the TCRαβ^+^ IEL subsets, we predicted a developmental relationship between memory-like and effector-like clusters. RNA velocity analysis of TCRαβ^+^ dataset displayed a clear trajectory originating from *Tcf7^+^* DN cells progressing toward CD8αα^+^ cells, with directional transition from memory-like to effector-like clusters, consistent with findings reported by Wang et. al. ^35^ . In contrast, the presence of two transcriptionally distinct DN subsets within the TCRγδ^+^ IELs may suggest the existence of two independent precursor populations: one *Tcf7^+^*subset giving rise to memory-like CD8αα^+^ cells, and a *Zeb2^+^* subset giving rise to effector-like populations. This model is supported by a recent study showing that TCRγδ^+^ CD8αα^+^ IELs from the colon of *Tcf7* knockout mice exhibited reduced *Cd160* expression but increased expression of effector and cytotoxic genes including *Il2rb*, *Gzma*, *Gzmb*, and *Tnfrsf9* ^73^. These findings suggest that effector-like cells can develop independently of memory-like cells likely originating from a distinct precursor lineage.

In summary, our study characterizes the transcriptional heterogeneity of TCRαβ^+^ and TCRγδ^+^ IELs, complementing previous finings and offering deeper insight into their complexity. We identified memory-like and effector-like subpopulations of IELs that exhibit strikingly similar transcriptional profiles between TCRαβ^+^ and TCRγδ^+^ lineages, despite their presumed participation in different immune responses. Moreover, our analyses suggest precursor-progeny relationships between DN and CD8αα^+^ cells, as well as between memory-like and effector-like CD8αα^+^ IELs, supporting a model in which these populations represent different stages of differentiation pathway. Together, our findings expand upon previously described IEL heterogeneity and underscore the need for further investigation into the functional relevance of transcriptionally similar TCRαβ^+^ and TCRγδ^+^ subsets.

## METHODS

### Ethics statement

This study was conducted in accordance with the Guide for Care and Use of Laboratory Animals of the National Institutes of Health. All animals were maintained and handled under approved Institutional Animal Care and Use Committee (IACUC) protocol (#5781) of the University of Massachusetts.

### Animals

C57BL/6J (stock no. 000664) mice were obtained from the Jackson Laboratory. All breedings were maintained at the University of Massachusetts, Amherst. This study was performed in accordance with the recommendations in the Guide for the Care and Use of Laboratory Animals of the National Institutes of Health. All animals were handled according to approved institutional animal care and use committee (IACUC) protocols of the University of Massachusetts.

### IEL isolation

The small intestine and colon were first removed from the rest of the gastrointestinal tract, onto collection media (RPMI supplemented with 25mM HEPES, 1% L-glutamine, 1% penicillin/streptomycin, 50µM β-mercaptoethanol, and 3% FBS). Peyer’s patches lining the small intestine were removed. Tissues were cleaned by flushing out feces with collection media and rinsing in PBS. Tissue fragments were agitated at 37°C for 20 minutes in collection media containing 5mM EDTA and 1mM DTT, then further shaken in serum-free collection media containing 2mM EDTA. The suspension was washed several times in collection media, and the IELs were collected as cells that passed through a 70µm filter. Finally, IELs were resuspended in collection media containing 10% FBS.

### Flow cytometry analysis

Flow cytometry data were acquired on BD LSR Fortessa. The following monoclonal antibodies from BioLegend were used: CD45.2 (104), CD45 (30-F11), TCRβ (H57-597), TCR8 (GL3), CD8α (53-6.7), CD8β (YTS156.7.7), Biotin-CD122 (5H3) and Streptavidin- AF647. The monoclonal CD4 (GK1.5) antibody and Brilliant Stain Buffer were obtained from BD Biosciences. The monoclonal CD160 (CNX46-3) antibody was obtained from eBiosciences.

Live cells were treated with anti-CD16/32 Fc block (2.4G2, BD Pharmingen) prior to staining with antibodies against surface markers. Staining for surface proteins was performed at 4 °C for 40 min, and FACS buffer (PBS + 0.5% BSA + 0.01% sodium azide) was used for washes. Data from BD LSR Fortessa were analyzed in FlowJo™ v10.9.0 Software. IEL populations were analyzed as shown in supplementary figure 5.

### Single-cell RNA sequencing

Cells were stained for CD45.2, TCRβ, TCR8, and sorted using the BD FACSAria Fusion instrument (BD Biosciences). After sorting, cells were counted using a Cellometer K2 cell counter (Nexcelom Bioscience) and by manual counting via hemocytometer. Single cell gene expression profiling was performed using the Chromium Next GEM Single Cell 3ʹ v3.1 (Dual Index) kit. Each cell suspension was loaded onto a well of Chip G on the 10x Genomics Chromium Controller System following the manufacturer’s user manual (10x Genomics). Barcoding and cDNA synthesis were performed according to the manufacturer’s instructions. Qualitative analysis of cDNA was performed using the 2100 Agilent Bioanalyzer High Sensitivity assay. The cDNA libraries were constructed using the 10x Chromium Single cell 3’ Library Kit v3.1 (dual index) according to the manufacturer’s protocol. Quality assessment of final libraries was done on Qubit fluorometer using a DNA High Sensitivity assay (Thermo Scientific) and a 2100 Agilent Bioanalyzer High Sensitivity assay (Agilent Technologies). Libraries were sequenced on an Illumina NextSeq 500 using the NextSeq 500/550 Mid Output Kit v2.5 (150 Cycles) sequencing kit, with the following read length: 28 bp Read1 for the 10x cell barcode and UMI, 90 bp Read 2 for the insert, and 10 bp I7 and I5 for the sample index. Phix (Illumina) was spiked in at 1% as per kit manual recommendation (10x Genomics).

The 10x Could Analysis Cell Ranger Pipeline (Cellranger version 7.1.0) was used to align reads and generate feature-barcode matrices. Reads were aligned to *Mus musculus reference genome* (Mouse GRCm39). The *aggr* pipeline was used to combine data from multiple samples into an experiment-wide feature-barcode matrix and analysis. The 10x Genomics Loupe Browser was used for visualization, initial quality assessment, and filtering of single cell gene expression data. Single Cell Gene Expression was performed at the Genomics Resource Laboratory, University of Massachusetts Amherst, MA.

### Analysis of scRNA-seq

Data analyses were performed using the Seurat package (version 4.3) in the R software version 4.2.1. The TCRαβ^+^ and TCRγδ^+^ datasets were analyzed individually but with identical procedures. The Seurat object was first generated by keeping all genes expressed by at least 3 cells. Cells were kept if they contained at least 100 unique features, at least 100 reads, and less than 10% mitochondrial genes. Data were normalized, scaled, then the top 2000 variable genes were used for the Principal Component Analysis (PCA) and generation of Uniform Manifold Approximation and Projection (UMAP) plots. Immune cells were filtered by expression of *Ptprc* > 0.5, then for T cells by keeping cells with at least 0.05% expression of *Cd3e*/*Cd3g*/*Cd3d*. Cluster identities were interpreted by calculating unique cluster marker genes and analyzing the distribution of known marker genes. Clusters with similar transcriptional profiles and localization on the UMAP plot were merged into groups. Poorly defined clusters were removed from further analysis.

Re-clustering of “CD8αα^+^” annotated cells from Figure 1 was performed by subsetting the “CD8αα^+^ *Tcf7*^+^”, “CD8αα^+^ *Prdm1*^+^”, and “*Ki67*^+^” clusters for the TCRαβ^+^ dataset, or the “CD8αα^+^ *Tcf7*^+^”, “CD8αα^+^ *Prdm1*^+^”, “CD8αα^+^ *Zeb2*^+^”, and “*Ki67*^+^” clusters for the TCRγδ^+^ dataset. Cells expressing *Cd4* or *Cd8b1* were removed, and the remaining cells were processed by the normalization and clustering steps as above. After examining the expression of *Cd8a*, cells were further separated into the CD8αα^+^ cells which expressed *Cd8a*, or DN cells which did not express *Cd8a*. Differentially expressed genes between clusters were calculated by the Wilcoxon ranked test. The list of transcription factors in the mouse genome were obtained from the FANTOM5 database. The most differentially expressed transcription factors between clusters were selected to show as bubble plots (**Fig. 2b, d and Fig. 3b, d**) by significance (adjusted p-value < 0.05) and proportion (difference in the percentage of cells expressing the gene > 20% for **Fig. 2b, d and Fig. 3b**, > 30% for **Fig. 3d**). RNA velocity analysis was performed using scVelo and Velocyto. Integration of the TCRαβ^+^ and TCRγδ^+^ datasets was performed after preliminary analysis of individual datasets with the SelectIntegrationFeatures(), FindIntegrationAnchors(), and IntegrateData() Seurat functions. The integrated dataset was preprocessed following standard methods, and clusters were defined at low resolution (0.1) to determine which cell types colocalize on the UMAP plot.

## Statistical analysis

Data statistical analysis was performed with Prism 9 (GraphPad software). *P*-values were determined using a two-tailed paired t-test, or one-way ANOVA.

## Data availability

The sequencing data that support the findings of this study has been deposited in the National Center of Biotechnology Information Gene Expression Omnibus (GEO) and is accessible through the accession number GSE284856.

## Acknowledgments

This work was supported by NIH grants AI146188 (L.A.P.), AI133041 (L.A.P.) and Biotechnology Training Program (BTP) of National Research Service Award T32 GM13096 (K.A.H. and A.C.L.).

## Author contributions

K.A.H., E.L.P. and L.A.P. designed the study. K.A.H., X.L., A.C.L., R.R. performed experiments. K.A.H. provided expertise on data analysis. E.L.P. and L.A.P. supervised the study. K.A.H., E.L.P. and L.A.P. wrote the manuscript.

## Competing interests

The authors declare no competing interests.

Correspondence and requests for materials should be addressed to Elena L. Pobezinskaya or Leonid A. Pobezinsky.

---

**Supplementary Figure 1:**
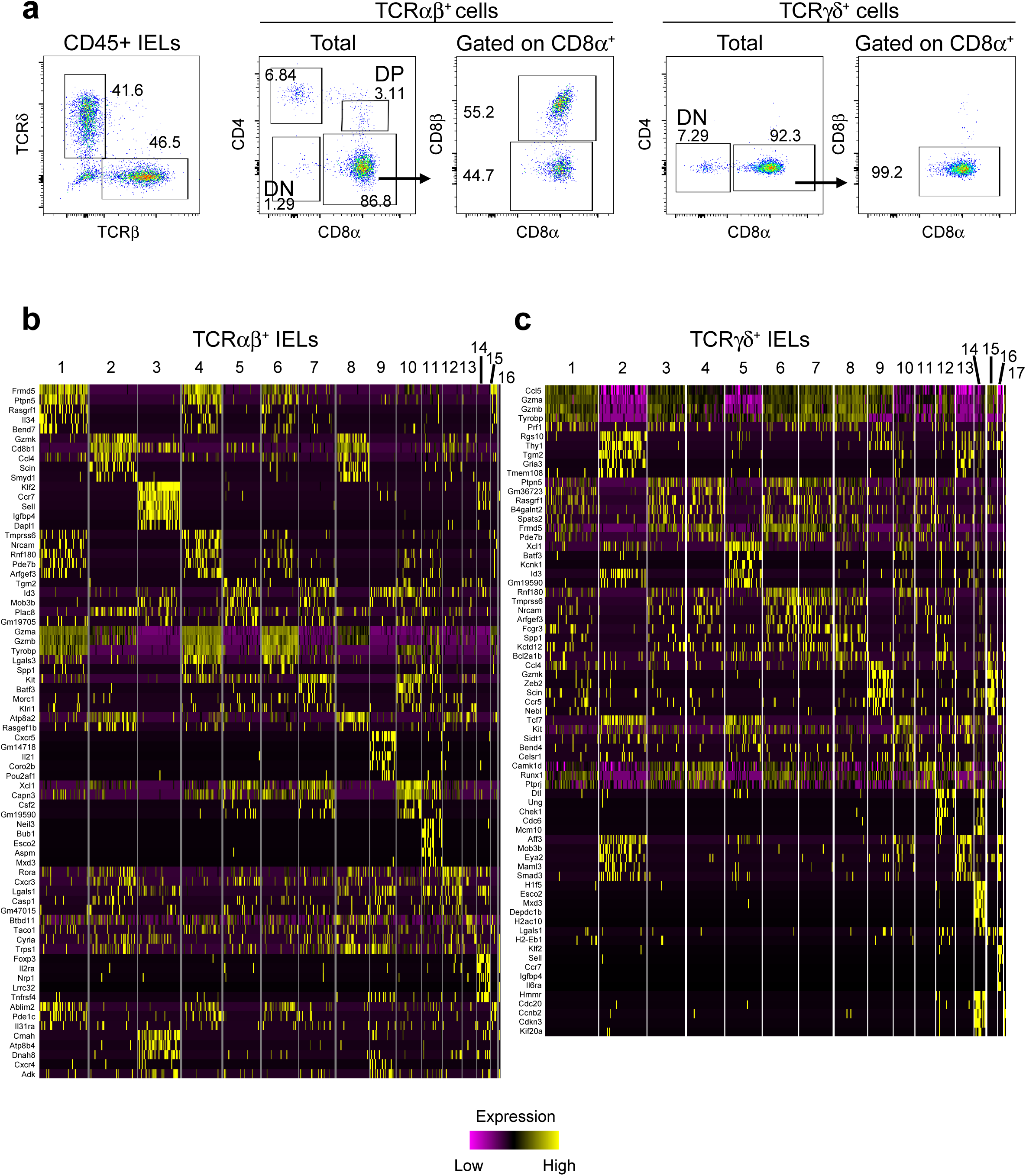
**a.** Flow cytometry plots representing the distribution of small intestinal IEL subpopulations among CD45^+^ cells. **b/c.** Heatmaps of cluster marker genes for the TCRαβ^+^ (a) and TCRγδ^+^ (b) datasets.

**Supplementary Figure 2:**
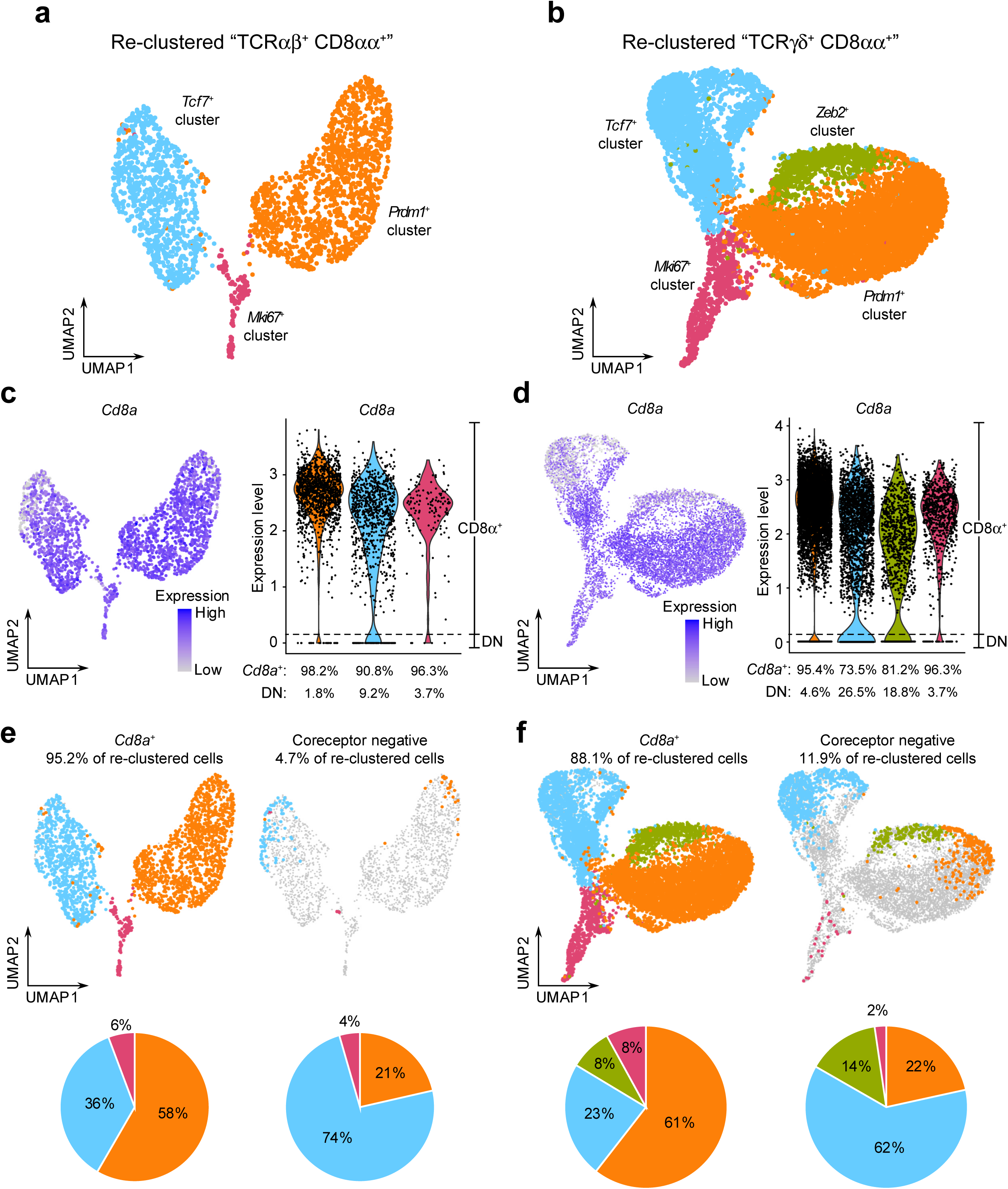
**a/b.** UMAP plots of re-clustered cells from the “CD8αα^+^” clusters in Figure 1 for the TCRαβ^+^ (a) and TCRγδ^+^ (b) datasets. Prior to re-clustering, cells expressing *Cd4* or *Cd8b1* coreceptor genes were removed. **c/d.** Expression of *Cd8a* in the re-clustered cells by visualizing the distribution on the UMAP plot and by violin plot for TCRαβ^+^ (c) and TCRγδ^+^ (d) datasets. **e/f.** Distribution of cell subsets either expressing the *Cd8a* coreceptor gene (CD8α^+^) or negative for all coreceptor genes (DN), and pie chart representation of annotated populations within each subset for TCRαβ^+^ (e) and TCRγδ^+^ (f) datasets.

**Supplementary Figure 3:**
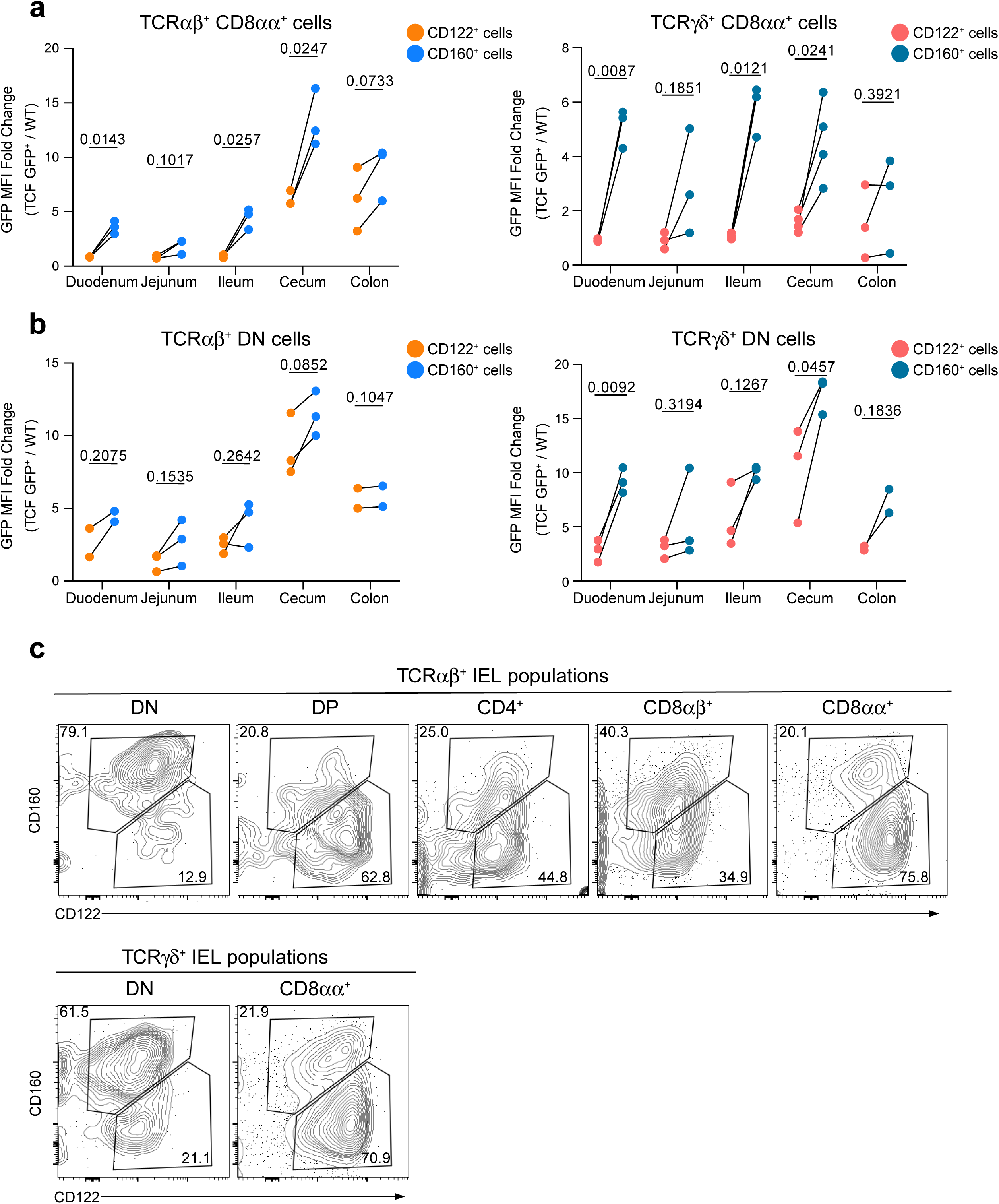
**a.** Quantification of GFP expression between TCRαβ^+^ (left) or TCRγδ^+^ (right) CD8αα^+^ CD122^int^CD160^+^ cells and CD122^hi^CD160^-^ cells from *Tcf7*-GFP mice, represented as median fluorescence intensity fold change compared to WT controls. **b.** Quantification of GFP expression between TCRαβ^+^ (left) or TCRγδ^+^ (right) DN CD122^int^CD160^+^ cells and CD122^hi^CD160^-^ cells from *Tcf7*-GFP mice, represented as median fluorescence intensity fold change compared to WT controls. **c.** Representative flow cytometry plots for the distribution of CD122 and CD160 among TCRαβ^+^ (a) or TCRγδ^+^ (b) IEL subpopulations. Data were analyzed by paired T-test (a, b).

**Supplementary Figure 4:**
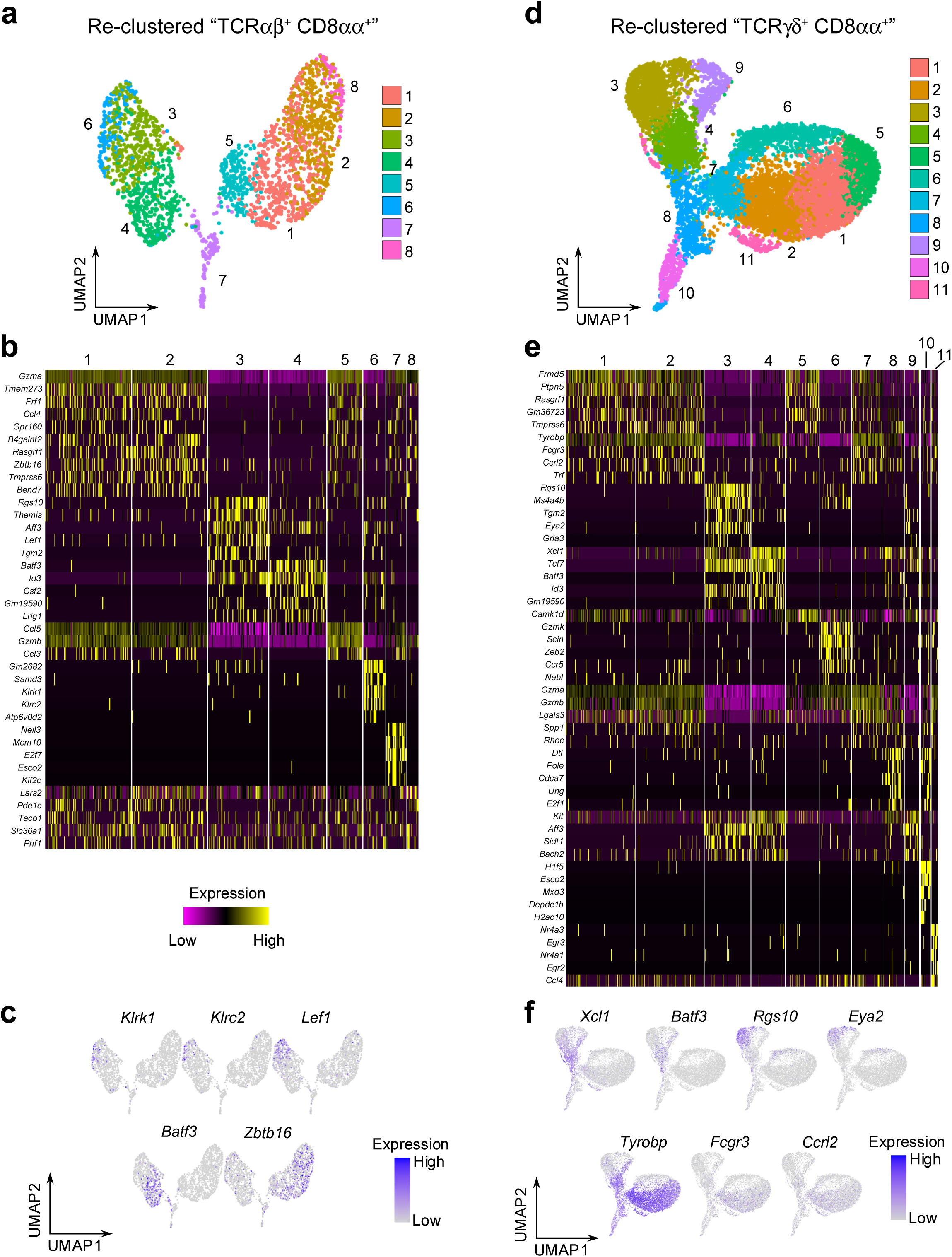
**a.** UMAP plots of re-clustered CD8αα^+^ and DN cells as in Supplementary Figure 2 with cluster number identities instead of phenotype annotations, for the TCRαβ^+^ dataset. **b.** Heatmaps of cluster marker genes for the re-clustered cells in the TCRαβ^+^ dataset. **c.** Representative UMAP plots for genes differentially expressed among re-clustered cells in the TCRαβ^+^ dataset. **d.** UMAP plots of re-clustered CD8αα^+^ and DN cells as in Supplementary Figure 2 with cluster number identities instead of phenotype annotations, for the TCRγδ^+^ dataset. **e.** Heatmaps of cluster marker genes for the re-clustered cells in the TCRγδ^+^ dataset. **f.** Representative UMAP plots for genes differentially expressed among re-clustered cells in the TCRγδ^+^ dataset.

**Supplementary Figure 5:**
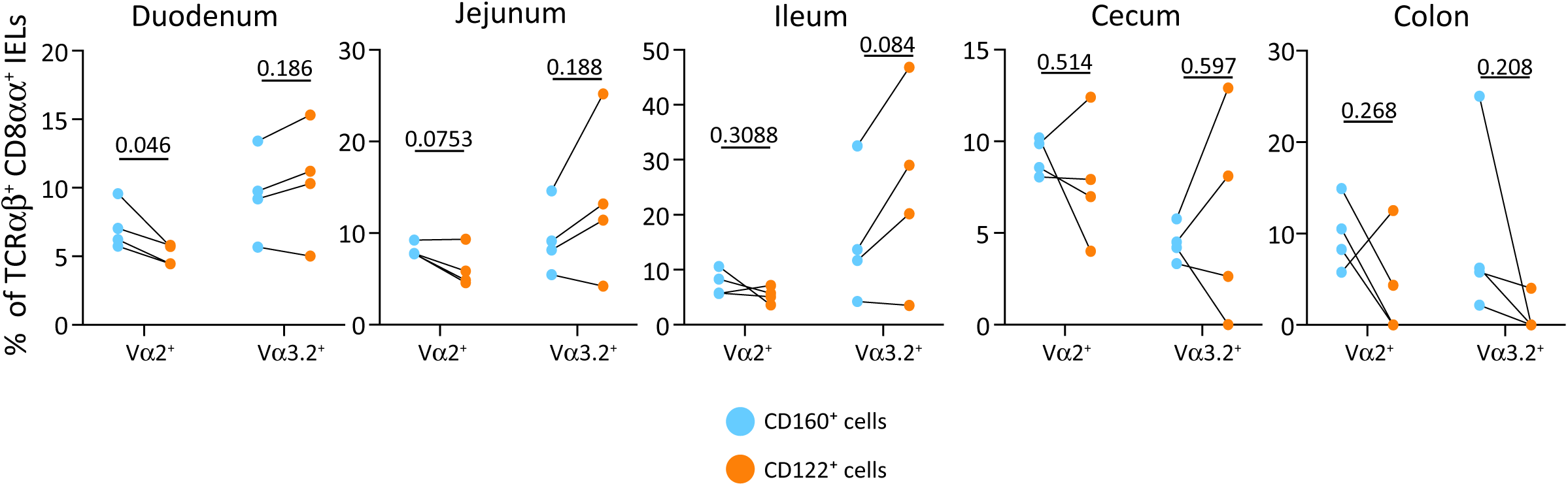
Percentage of TCRαβ^+^ CD8αα^+^ CD122^int^CD160^+^ or CD122^hi^CD160^-^ IELs expressing the TCRVα2 or TCRVα3.2 chain. Data were analyzed by Paired T-test.

**Supplementary Figure 6:**
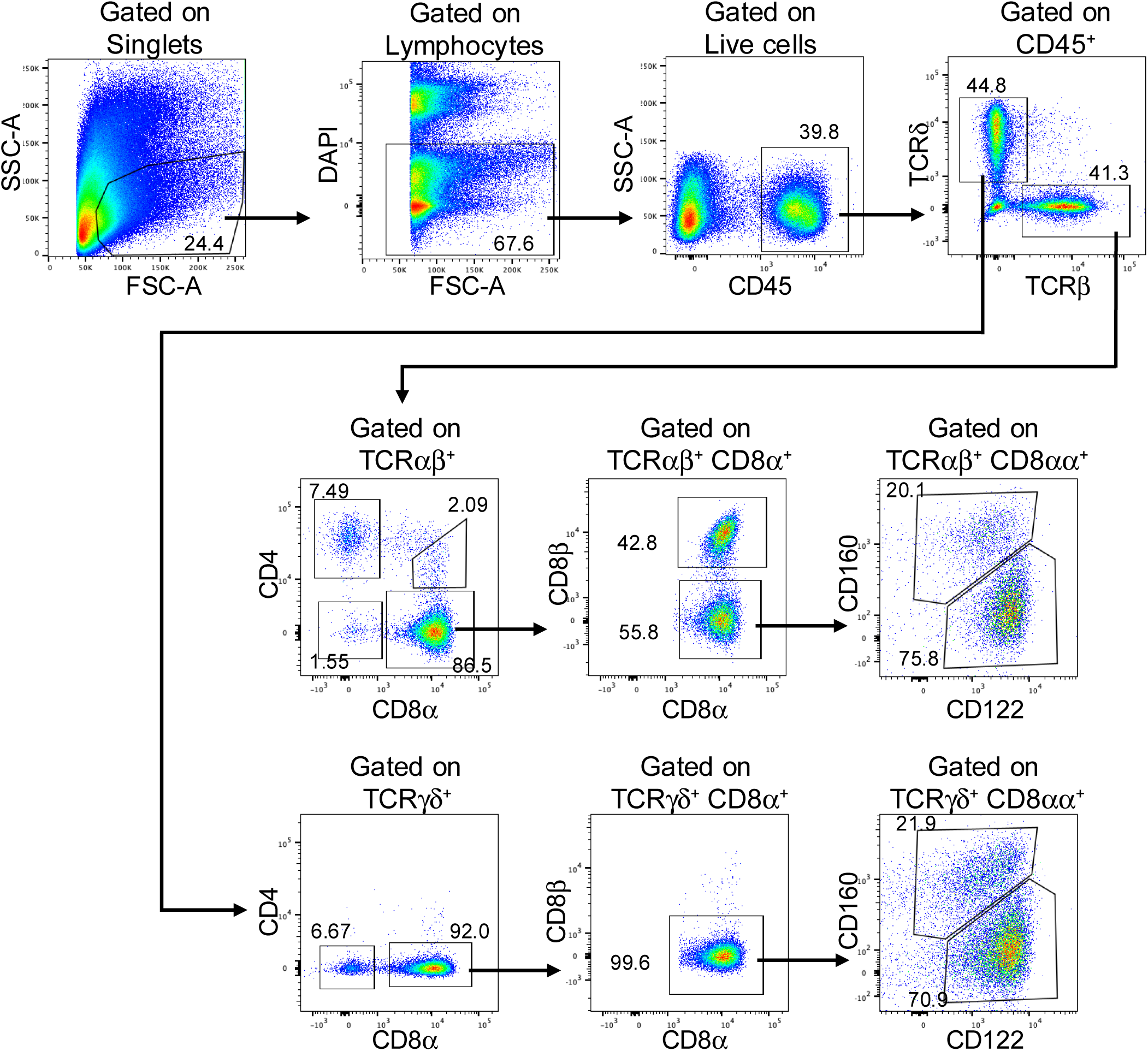
Representative flow cytometry gating scheme used for IEL experiments.

## References

1. Chen, Y., Chou, K., Fuchs, E., Havran, W.L. & Boismenu, R. Protection of the intestinal mucosa by intraepithelial gamma delta T cells. Proc Natl Acad Sci U S A 99, 14338–43 (2002).

2. Cheroutre, H., Lambolez, F. & Mucida, D. The light and dark sides of intestinal intraepithelial lymphocytes. Nat Rev Immunol 11, 445–56 (2011).

3. Han, J. et al. TGF-beta controls development of TCRgammadelta(+)CD8alphaalpha(+) intestinal intraepithelial lymphocytes. Cell Discov 9, 52 (2023).

4. Olivares-Villagomez, D. & Van Kaer, L. Intestinal Intraepithelial Lymphocytes: Sentinels of the Mucosal Barrier. Trends Immunol 39, 264–275 (2018).

5. Roberts, S.J. et al. T-cell alpha beta + and gamma delta + deficient mice display abnormal but distinct phenotypes toward a natural, widespread infection of the intestinal epithelium. Proc Natl Acad Sci U S A 93, 11774–9 (1996).

6. Ma, H., Qiu, Y. & Yang, H. Intestinal intraepithelial lymphocytes: Maintainers of intestinal immune tolerance and regulators of intestinal immunity. J Leukoc Biol 109, 339–347 (2021).

7. Guy-Grand, D. et al. Two gut intraepithelial CD8+ lymphocyte populations with different T cell receptors: a role for the gut epithelium in T cell differentiation. J Exp Med 173, 471–81 (1991).

8. Huang, Y. et al. Mucosal memory CD8(+) T cells are selected in the periphery by an MHC class I molecule. Nat Immunol 12, 1086–95 (2011).

9. Masopust, D., Vezys, V., Marzo, A.L. & Lefrancois, L. Preferential localization of effector memory cells in nonlymphoid tissue. Science 291, 2413–7 (2001).

10. Mucida, D. et al. Transcriptional reprogramming of mature CD4(+) helper T cells generates distinct MHC class II-restricted cytotoxic T lymphocytes. Nat Immunol 14, 281–9 (2013).

11. Shires, J., Theodoridis, E. & Hayday, A.C. Biological insights into TCRgammadelta+ and TCRalphabeta+ intraepithelial lymphocytes provided by serial analysis of gene expression (SAGE). Immunity 15, 419–34 (2001).

12. Das, G. et al. An important regulatory role for CD4+CD8 alpha alpha T cells in the intestinal epithelial layer in the prevention of inflammatory bowel disease. Proc Natl Acad Sci U S A 100, 5324–9 (2003).

13. McDonald, B.D., Jabri, B. & Bendelac, A. Diverse developmental pathways of intestinal intraepithelial lymphocytes. Nat Rev Immunol 18, 514–525 (2018).

14. Groux, H. et al. A CD4+ T-cell subset inhibits antigen-specific T-cell responses and prevents colitis. Nature 389, 737–42 (1997).

15. Maynard, C.L. et al. Regulatory T cells expressing interleukin 10 develop from Foxp3+ and Foxp3- precursor cells in the absence of interleukin 10. Nat Immunol 8, 931–41 (2007).

16. Sujino, T. et al. Tissue adaptation of regulatory and intraepithelial CD4(+) T cells controls gut inflammation. Science 352, 1581–6 (2016).

17. Cervantes-Barragan, L. et al. Lactobacillus reuteri induces gut intraepithelial CD4(+)CD8alphaalpha(+) T cells. Science 357, 806–810 (2017).

18. Hoytema van Konijnenburg, D.P., et al. Intestinal Epithelial and Intraepithelial T Cell Crosstalk Mediates a Dynamic Response to Infection. Cell 171, 783–794 e13 (2017).

19. Wojciech, L. et al. Non-canonicaly recruited TCRalphabetaCD8alphaalpha IELs recognize microbial antigens. Sci Rep 8, 10848 (2018).

20. Kornberg, A., et al. Gluten induces rapid reprogramming of natural memory alphabeta and gammadelta intraepithelial T cells to induce cytotoxicity in celiac disease. Sci Immunol 8, eadf4312 (2023).

21. Pobezinsky, L.A. et al. Clonal deletion and the fate of autoreactive thymocytes that survive negative selection. Nat Immunol 13, 569–78 (2012).

22. Denning, T.L. et al. Mouse TCRalphabeta+CD8alphaalpha intraepithelial lymphocytes express genes that down-regulate their antigen reactivity and suppress immune responses. J Immunol 178, 4230–9 (2007).

23. Guehler, S.R., Finch, R.J., Bluestone, J.A. & Barrett, T.A. Increased threshold for TCR-mediated signaling controls self reactivity of intraepithelial lymphocytes. J Immunol 160, 5341–6 (1998).

24. Kurd, N.S. et al. Factors that influence the thymic selection of CD8alphaalpha intraepithelial lymphocytes. Mucosal Immunol 14, 68–79 (2021).

25. Tiberti, S. et al. GZMK(high) CD8(+) T effector memory cells are associated with CD15(high) neutrophil abundance in non-metastatic colorectal tumors and predict poor clinical outcome. Nat Commun 13, 6752 (2022).

26. Yomogida, K. et al. The transcription factor Aiolos restrains the activation of intestinal intraepithelial lymphocytes. Nat Immunol 25, 77–87 (2024).

27. Boismenu, R. & Havran, W.L. Modulation of epithelial cell growth by intraepithelial gamma delta T cells. Science 266, 1253–5 (1994).

28. Inagaki-Ohara, K. et al. Mucosal T cells bearing TCRgammadelta play a protective role in intestinal inflammation. J Immunol 173, 1390–8 (2004).

29. Komano, H. et al. Homeostatic regulation of intestinal epithelia by intraepithelial gamma delta T cells. Proc Natl Acad Sci U S A 92, 6147–51 (1995).

30. Kuhl, A.A. et al. Aggravation of intestinal inflammation by depletion/deficiency of gammadelta T cells in different types of IBD animal models. J Leukoc Biol 81, 168–75 (2007).

31. Jordan-Paiz, A. et al. CXCR5(+)PD-1(++) CD4(+) T cells colonize infant intestines early in life and promote B cell maturation. Cell Mol Immunol 20, 201–213 (2023).

32. Leonard, W.J., Lin, J.X. & O’Shea, J.J. The gamma(c) Family of Cytokines: Basic Biology to Therapeutic Ramifications. Immunity 50, 832–850 (2019).

33. Lynch, A., et al. A Dapl1+ subpopulation of naïve CD8 T cells contains committed precursors of memory lineage. bioRxiv (2025).

34. Nguyen, Q.P. et al. Transcriptional programming of CD4(+) T(RM) differentiation in viral infection balances effector- and memory-associated gene expression. Sci Immunol 8, eabq7486 (2023).

35. Wang, Y.C. et al. Intestinal cell type-specific communication networks underlie homeostasis and response to Western diet. J Exp Med 220 (2023).

36. Panda, S.K. et al. Repression of the aryl-hydrocarbon receptor prevents oxidative stress and ferroptosis of intestinal intraepithelial lymphocytes. Immunity 56, 797–812 e4 (2023).

37. Huang, J. et al. Intraepithelial lymphocytes promote intestinal regeneration through CD160/HVEM signaling. Mucosal Immunol 17, 257–271 (2024).

38. Taveirne, S. et al. Inhibitory receptors specific for MHC class I educate murine NK cells but not CD8alphaalpha intestinal intraepithelial T lymphocytes. Blood 118, 339–47 (2011).

39. James, O.J. et al. IL-15 and PIM kinases direct the metabolic programming of intestinal intraepithelial lymphocytes. Nat Commun 12, 4290 (2021).

40. Lee, G.A. & Liao, N.S. CD8(+)CD122(+) T cell homeostasis is controlled by different levels of IL-15 trans-presentation. J Microbiol Immunol Infect 54, 514–517 (2021).

41. Ma, L.J., Acero, L.F., Zal, T. & Schluns, K.S. Trans-presentation of IL-15 by intestinal epithelial cells drives development of CD8alphaalpha IELs. J Immunol 183, 1044–54 (2009).

42. Tan, C.L. et al. CD160 Stimulates CD8(+) T Cell Responses and Is Required for Optimal Protective Immunity to Listeria monocytogenes. Immunohorizons 2, 238–250 (2018).

43. Beagley, K.W. et al. Differences in intraepithelial lymphocyte T cell subsets isolated from murine small versus large intestine. J Immunol 154, 5611–9 (1995).

44. Boll, G., Rudolphi, A., Spiess, S. & Reimann, J. Regional specialization of intraepithelial T cells in the murine small and large intestine. Scand J Immunol 41, 103–13 (1995).

45. Chen, B. et al. Commensal Bacteria-Dependent CD8alphabeta(+) T Cells in the Intestinal Epithelium Produce Antimicrobial Peptides. Front Immunol 9, 1065 (2018).

46. Hung, C.T. et al. Western diet reduces small intestinal intraepithelial lymphocytes via FXR-Interferon pathway. Mucosal Immunol 17, 1019–1028 (2024).

47. Park, C. et al. Obesity Modulates Intestinal Intraepithelial T Cell Persistence, CD103 and CCR9 Expression, and Outcome in Dextran Sulfate Sodium-Induced Colitis. J Immunol 203, 3427–3435 (2019).

48. Rodriguez-Marino, N. et al. Dietary fiber promotes antigen presentation on intestinal epithelial cells and development of small intestinal CD4(+)CD8alphaalpha(+) intraepithelial T cells. Mucosal Immunol 17, 1301–1313 (2024).

49. Yang, Q. et al. TCF-1 upregulation identifies early innate lymphoid progenitors in the bone marrow. Nat Immunol 16, 1044–50 (2015).

50. Chang, J.W., Koike, T. & Iwashima, M. hnRNP-K is a nuclear target of TCR- activated ERK and required for T-cell late activation. Int Immunol 21, 1351–61 (2009).

51. Gallardo, M. et al. hnRNP K Is a Haploinsufficient Tumor Suppressor that Regulates Proliferation and Differentiation Programs in Hematologic Malignancies. Cancer Cell 28, 486–499 (2015).

52. Li, H. et al. YBX1 as an oncogenic factor in T-cell acute lymphoblastic leukemia. Blood Adv 7, 4874–4885 (2023).

53. Gui, Y. et al. Development and function of natural TCR(+) CD8alphaalpha(+) intraepithelial lymphocytes. Front Immunol 13, 1059042 (2022).

54. Cao, M.Y., Davidson, D., Yu, J., Latour, S. & Veillette, A. Clnk, a novel SLP-76- related adaptor molecule expressed in cytokine-stimulated hemopoietic cells. J Exp Med 190, 1527–34 (1999).

55. Chou, C. et al. Programme of self-reactive innate-like T cell-mediated cancer immunity. Nature 605, 139–145 (2022).

56. Lehto, M. et al. Targeting of OSBP-related protein 3 (ORP3) to endoplasmic reticulum and plasma membrane is controlled by multiple determinants. Exp Cell Res 310, 445–62 (2005).

57. McNerney, M.E., Lee, K.M. & Kumar, V. 2B4 (CD244) is a non-MHC binding receptor with multiple functions on natural killer cells and CD8+ T cells. Mol Immunol 42, 489–94 (2005).

58. Wang, T. et al. FERM-containing protein FRMD5 is a p120-catenin interacting protein that regulates tumor progression. FEBS Lett 586, 3044–50 (2012).

59. Yan, J. et al. FGL2 promotes tumor progression in the CNS by suppressing CD103(+) dendritic cell differentiation. Nat Commun 10, 448 (2019).

60. Koh, J.Y., Kim, D.U., Moon, B.H. & Shin, E.C. Human CD8(+) T-Cell Populations That Express Natural Killer Receptors. Immune Netw 23, e8 (2023).

61. Wensveen, F.M., Jelencic, V. & Polic, B. NKG2D: A Master Regulator of Immune Cell Responsiveness. Front Immunol 9, 441 (2018).

62. Savage, A.K. et al. The transcription factor PLZF directs the effector program of the NKT cell lineage. Immunity 29, 391–403 (2008).

63. Zhang, S., Laouar, A., Denzin, L.K. & Sant’Angelo, D.B. Zbtb16 (PLZF) is stably suppressed and not inducible in non-innate T cells via T cell receptor-mediated signaling. Sci Rep 5, 12113 (2015).

64. Garcia-Bernal, D. et al. RGS10 restricts upregulation by chemokines of T cell adhesion mediated by alpha4beta1 and alphaLbeta2 integrins. J Immunol 187, 1264–72 (2011).

65. Zhang, T., Xu, J. & Xu, P.X. Eya2 expression during mouse embryonic development revealed by Eya2(lacZ) knockin reporter and homozygous mice show mild hearing loss. Dev Dyn 250, 1450–1462 (2021).

66. Hummel, J.F. et al. Single-cell RNA-sequencing identifies the developmental trajectory of C-Myc-dependent NK1.1(-) T-bet(+) intraepithelial lymphocyte precursors. Mucosal Immunol 13, 257–270 (2020).

67. Ruscher, R., Kummer, R.L., Lee, Y.J., Jameson, S.C. & Hogquist, K.A. CD8alphaalpha intraepithelial lymphocytes arise from two main thymic precursors. Nat Immunol 18, 771–779 (2017).

68. Ruscher, R. et al. Intestinal CD8alphaalpha IELs derived from two distinct thymic precursors have staggered ontogeny. J Exp Med 217(2020).

69. Jaeger, N. et al. Single-cell analyses of Crohn’s disease tissues reveal intestinal intraepithelial T cells heterogeneity and altered subset distributions. Nat Commun 12, 1921 (2021).

70. Yonemoto, Y. et al. Single cell analysis revealed that two distinct, unique CD4(+) T cell subsets were increased in the small intestinal intraepithelial lymphocytes of aged mice. Front Immunol 15, 1340048 (2024).

71. Kurd, N.S., et al. Early precursors and molecular determinants of tissue-resident memory CD8(+) T lymphocytes revealed by single-cell RNA sequencing. Sci Immunol 5(2020).

72. Milner, J.J. et al. Heterogenous Populations of Tissue-Resident CD8(+) T Cells Are Generated in Response to Infection and Malignancy. Immunity 52, 808–824 e7 (2020).

73. Yakou, M.H., et al. TCF-1 limits intraepithelial lymphocyte antitumor immunity in colorectal carcinoma. Sci Immunol 8, eadf2163 (2023).

74. Luangsay, S. et al. CCR5 mediates specific migration of Toxoplasma gondii- primed CD8 lymphocytes to inflammatory intestinal epithelial cells. Gastroenterology 125, 491–500 (2003).

75. Olive, A.J., Gondek, D.C. & Starnbach, M.N. CXCR3 and CCR5 are both required for T cell-mediated protection against C. trachomatis infection in the murine genital mucosa. Mucosal Immunol 4, 208–16 (2011).

76. Omilusik, K.D. et al. Transcriptional repressor ZEB2 promotes terminal differentiation of CD8+ effector and memory T cell populations during infection. J Exp Med 212, 2027–39 (2015).

